# How do stochastic processes and genetic threshold effects explain incomplete penetrance and inform causal disease mechanisms?

**DOI:** 10.1101/2023.10.04.560918

**Authors:** Dagan Jenkins

## Abstract

Incomplete penetrance is the rule rather than the exception in Mendelian disease. In syndromic monogenic disorders, phenotypic variability can be viewed as the combination of incomplete penetrance for each of multiple independent clinical features. Within genetically identical individuals, such as isogenic model organisms, stochastic variation at molecular and cellular levels is the primary cause of incomplete penetrance according to a genetic threshold model. By defining specific probability distributions of causal biological readouts and genetic liability values, stochasticity and incomplete penetrance provide information about threshold values in biological systems. Ascertainment of threshold values has been achieved by simultaneous scoring of relatively simple phenotypes and quantitation of molecular readouts at the level of single cells. However, this is much more challenging for complex morphological phenotypes using experimental and reductionist approaches alone, where cause and effect are separated temporally and across multiple biological modes and scales. Here I consider how causal inference, which integrates observational data with high confidence causal models, might be used to quantify the relative contribution of different sources of stochastic variation to phenotypic diversity. Collectively, these approaches could inform disease mechanisms, improve predictions of clinical outcomes and prioritise gene therapy targets across modes and scales of gene function.

## Introduction

The identification of thousands of *bona fide* Mendelian disease genes represents one of the most important achievements in biomedical research, providing novel insights into disease mechanisms and promising to inform treatments and disease outcomes for patients [1–4]. However, the extensive phenotypic variability observed in patients with mutations in the same gene – even in pairs of monozygotic twins – reduces this predictive power and has proven to be a major limiting factor [5–9]. Given that most treatments represent the amelioration of severe forms of disease rather than being curative *per se*, it may be possible to identify new therapeutic targets by studying the mechanisms that underlie this variability and even to tailor medicines to individuals. Such work will also provide insight into fundamental genetic principles in the broad sense, or, as Waddington called it, *The Strategy of the Genes* [64]. This includes insight into the role of stochastic processes in biological systems, which is the focus of this Perspective.

Embryonic development and adult tissue homeostasis are exquisitely robust when viewed at macroscopic levels. As Wolpert discussed, it is bewildering that, for most people, their left and right arms remain almost identical in length and overall appearance throughout their lifetime, even though they have had no means of communicating with one another ever since the midline barrier was laid down in the early foetus [10]. This maintenance of mirror-image symmetry is also a feature of many other bilateral tissues. Yet when quantitative molecular and cellular assays are used to observe biological systems at microscopic levels they can be seen to be inherently noisy [11,12].

What then are the molecular and cellular sources of this noise? Stochastic processes can generate variation at the level of single cells, including intrinsic noise that results from non-statistical sampling of relatively small numbers of molecules. (For example, there are only two copies of each gene promoter located on an autosome). This is thought to arise from processes such as transcriptional and translational bursting events [11–16] and can be ascertained by assessment of cell-to-cell variation.

By contrast extrinsic noise may arise from environmental fluctuations, and variability in the response to the environment may result from the disparity of cell state [17–20]. We might expect that extrinsic environmental factors such as temperature or diet are likely to be uniform across a tissue and generate inter-individual variation rather than noise at the single-cell level. Therefore, different sources of stochastic variation may have different signatures at the single molecule, cellular and tissue/inter-individual levels.

At the transcriptional level, the relative contributions of intrinsic versus extrinsic noise can be evaluated according to the correlation of pre-processed transcripts present at two identical promoters in single cells, such as the two alleles of an autosomal gene in an isogenic line [11,17,19]. Intrinsic vs. extrinsic noise is indicated by the degree of allelic imbalance. Stochastic processes of this type have been demonstrated in a number of systems, and they have been shown to underlie incomplete penetrance, as will be outlined in detail below.

## What is phenotypic variability?

Many Mendelian traits are discrete phenotypes that segregate with a defined mutation within a family. Variation in the severity of these phenotypes may involve individuals who carry the mutation but are unaffected, which is defined as incomplete penetrance. Alternatively, affected individuals may vary in severity, known as variable expressivity.

Most Mendelian diseases are syndromes which are compound phenotypes involving the combination of multiple independent features affecting a variety of organs or tissues [4,21-23]. While these diseases do appear to segregate according to Mendel’s laws when considered as a whole, relatively few such disorders would actually exist without flexible systems of diagnosis [21–26]. For example, patients with Bardet-Biedl syndrome exhibit a combination of 6 major and 8 minor clinical features, and a clinical diagnosis of BBS is accepted only if a patient has four major criteria, or three major criteria and two or more secondary criteria [27,28]. For a series of genes causing brachydactylies, such as Robinow syndrome, heterozygous mutations cause isolated non-syndromic shortening of the digits while homozygous mutations in the same gene cause the same phenotype together with other syndromic features [29]. What seems to be the rule rather than the exception is that for these syndromes most individual clinical features are associated with incomplete penetrance when considered in isolation. For each independent phenotype mutations in Mendelian disease genes can be viewed as having three key features such that they are:

1. rare susceptibility alleles, with;
2. moderate to high penetrance, and;
3. pleiotropic effects.

We may therefore consider that clinical variability in syndromic Mendelian disorders reflects incomplete penetrance combined across a number of independent clinical features. Incomplete penetrance may also explain variable expressivity where higher-level phenotypes involve multiple repeating structures or result from the combination of several constituent endophenotypes across biological scales. As a hypothetical example, a quantitative trait such as glomerular filtration rate may be the product of incomplete penetrance for discrete effects on each of ∼10^6^ nephrons within the kidney. Alternatively, a complex limb malformation might reflect incomplete penetrance for multiple independent developmental processes regulating digit number, identity and joint formation, which are temporally integrated [30].

Within naturally breeding populations, such as humans with genetic diseases, incomplete penetrance and variable expressivity are attributed to modifiers owing to genetic and environmental heterogeneity amongst these individuals. Stochastic processes also contribute to clinical variability. In genetically identical experimental organisms in controlled environments, stochastic processes are the major source of variation. How stochasticity generates phenotypic variability is poorly understood, especially for complex morphological traits. As such, the 6,000 or so Mendelian disease genes that have been identified [4], and the series of mutations found within them, provide us with many models to investigate the role of stochastic processes in disease susceptibility.

### Incomplete penetrance and genetic threshold effects: stochastic variation informs threshold values for incompletely penetrant traits

The genetic threshold model was first proposed in 1934 by Wright [31,32], to explain the appearance of discrete phenotypes. It was invoked to explain preaxial polydactyly that was observed with different penetrance values in a number of strains of guinea pigs. There are several features of this model and historical aspects to consider. In its original form, the model was used to explain this discrete morphological trait in inbred (isogenic) lines of guinea pigs. As for polydactyly, a wide variety of phenotypes have subsequently been observed at background levels in isogenic lines of rodents, with strain-specific penetrance values [33–35]. In this early era of genetics and developmental biology (or embryology, as it was then known), Wright especially attributed this phenotypic variability to environmental variation. He considered that genetic background predisposed different strains to disease, and that these effects were modified by environmental factors. He was also open to the idea of stochasticity, although he didn’t use this term. Instead, he commented on, ‘…irregularities in development due to the intangible sort of causes to which the word chance is applied’ [36]. Wright particularly considered position effects within the uterus and differences in foetal blood supply as forms of environmental variation which are quite different from stochastic molecular processes.

The combination of all genetic and environmental modifiers impacting a particular phenotype in an individual is defined as the liability. It is important to realise that liability is an intangible construct that can’t be measured. It is not possible to define all genetic modifiers [37], not least because of statistical power considerations and small effect sizes [38,39], and the sum total of environmental factors are too complicated to realistically ascertain. By studying isogenic lines, Wright paved the way to remove genetic modifiers as a source of variation and greatly simplified the problem. This provided the basis for an experimentally tractable approach. We can now use gene-editing to introduce virtually any mutation in isogenic lines to study defined genetic variation in the context of a variety of fixed genetic backgrounds. We can therefore take essentially the same approach as Wright albeit in a molecularly targeted way.

A central difficulty for the theory of evolution was to understand the connection between particulate inheritance and quantitative phenotypic variation. In seminal work, Fisher demonstrated that quantitative traits could arise from particulate inheritance of multiple susceptibility alleles with small effect sizes across many different loci i.e. oligogenic traits. In this model, the liability in populations of genetically heterogenous individuals in varied environments is taken to be normally distributed [40–42]. Later, and presumably taking inspiration from Wright’s work, Cohen, in his consideration of the inheritance of pyloric stenosis was the first to combine this concept with the threshold model such that individuals carrying a greater number of modifiers than a threshold liability value exhibit the trait [66,67]. Falconer subsequently did much to extend this model and understand its implications [65]. This is a model that permeates genetics because it helps us to think about the intersection between genes and environment, quantitative variation and discrete phenotypes i.e. diseases. But by being based on the concept of liability it is not particularly useful.

While Fisher’s work was ground-breaking for evolutionary theory, its application in complex genetics placed the emphasis on heterogenous populations and oligogenic traits which forms the basis for the genetic threshold model typically seen in textbooks. This side-lined Wright’s approach which had focused on isogenic models as experimentally tractable systems. Conscious of this, we can now build on Wright’s model in the post-genomic era by considering the distribution of liability values for a *defined genotype* on a *specific genetic background* (with genotype referring to the Mendelian trait locus). In this situation, stochastic processes account for most of the variation in liability values. We must therefore *redefine liability to include naturally occurring molecular and cellular variation*. In this form, liability constitutes the functional effects of experimentally controlled genetic and environmental factors as well as stochastic variation in the form of tangible readouts such as gene product abundance (e.g. RNA and protein abundance) or activity (e.g. transcription factor binding to a promoter or a signalling output such as protein phosphorylation). The key is that such readouts can be measured, at least in principle. At the level of individuals, the liability value would fall below the threshold for unaffected individuals and above this threshold for those individuals that are affected. As will be discussed in the next section, these are not only readouts in the sense that is widely used in biomedical research, such as luciferase reporter assays or other bioreporters that are surrogate correlates of gene activity. They must be a node or combination of nodes on the causal path linking genotype to phenotype and capture the total stochastic variation within the wider network that impacts upon the phenotype (see below for a consideration of how this might be practically implemented using current technologies).

This leads us to several important features of the molecularly targeted genetic threshold model. Firstly, incomplete penetrance is the result of stochastic variation in liability values under experimentally controlled conditions for isogenic models - without this variation, all individuals would either be affected or unaffected. Incomplete penetrance means that the threshold liability value is located within the range of stochastic variation for a defined genotype in a particular strain. As we shall see in the next section, the penetrance value is exactly equal to the area under this curve that falls above the threshold liability value.

In the case of incomplete penetrance, stochastic variation therefore carries information about the threshold value in the form of its probability distribution function and the exact penetrance value for a particular trait.

### Threshold effects demonstrated by simultaneous assessment of phenotype and readout in single cells

A genetic threshold is an exact value that relates phenotype penetrance to the distribution of liability values for a specified genotype [Figure 1]. Where the liability values for a series of genotypes on an isogenic background are described by the same type of statistical distribution (Normal, Poisson, Gamma, etc) with the same variance, differing only in their population means, the penetrance values across a series of genotypes will be described by the cumulative distribution function (CDF) of liability values [Figure 1C]. (Note, this is an idealised model for illustration. Empirically, mutations are typically found to be associated with greater stochastic variation. Furthermore, different mutations may be associated with different variances according to their mechanism of pathogenesis and different buffering mechanisms that may modulate their effects (splicing, nonsense-mediated mRNA decay, chaperones/protein folding etc.). Nonetheless, phenotype penetrance is given by the CDF for a particular genotype if the appropriate readout(s) is ascertained).

**Figure 1.**
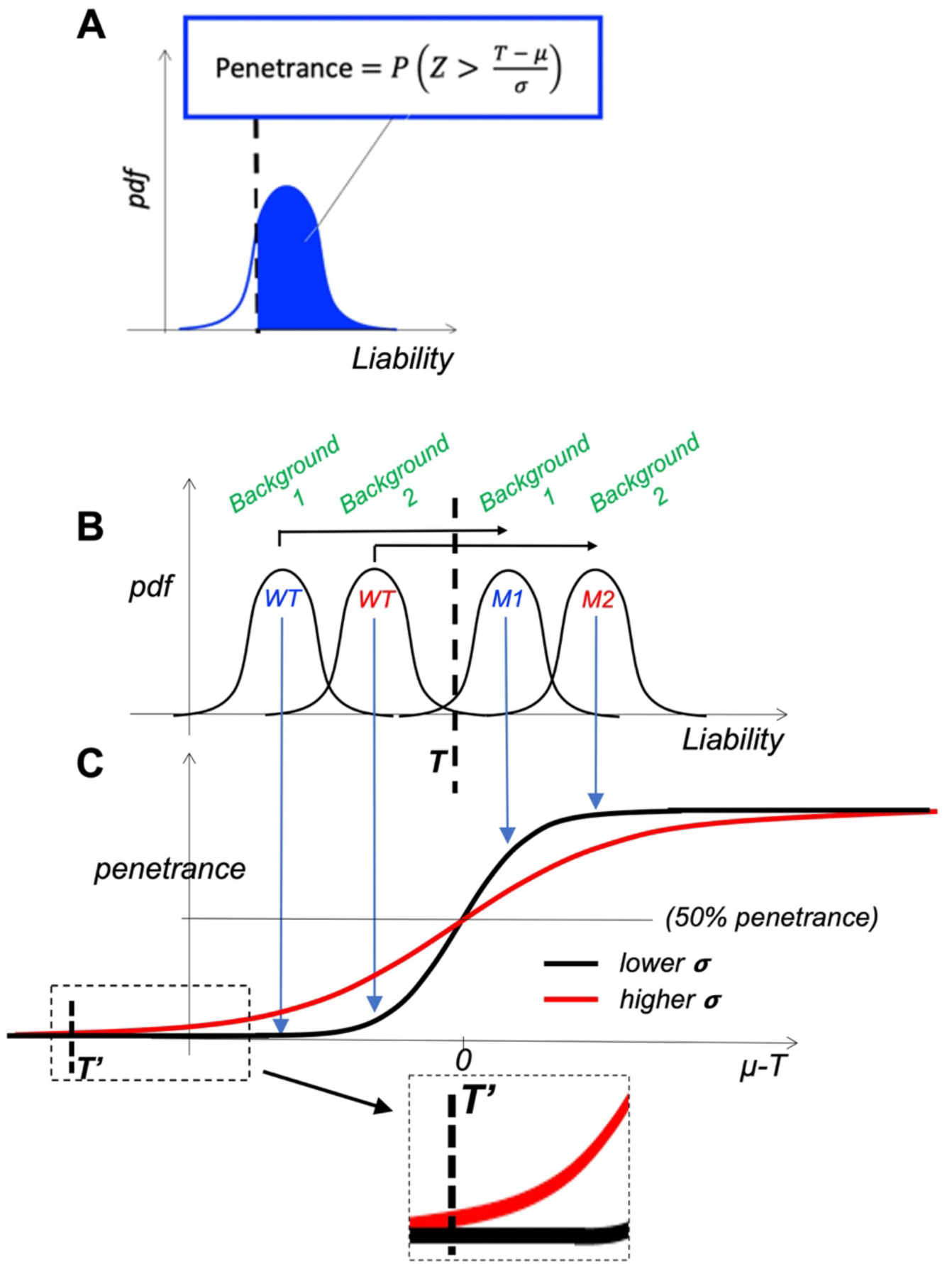
Genetic threshold model. **A.** Basic model showing how the penetrance value for a trait is directly derived from the probability distribution function (pdf) of liability values in relation to a genetic threshold value – incomplete penetrance occurs when the threshold value falls within the pdf associated with a mutation. **B.** Cartoon of pdfs for different mutations (M1, M2) on different genetic backgrounds showing that liability is the result of genetic background, mutational effect and stochastic variation. Different penetrance values for different genotypes are defined exactly by these pdfs in relation to the genetic threshold value (T, dashed line). **C.** Disease penetrance relates to mean liability values for pdfs defined by genetic background, mutation and stochastic processes in **B** according to cumulative distribution functions (black line). Red line illustrates the cumulative distribution function for the same set of distributions in **B**, but with higher levels of stochastic variation (note where the mean equals the threshold value, penetrance is 50% regardless of variance). The threshold value that can be ascertained by simultaneous ascertainment in single cells (T’) is a function of the genetic threshold value (T). Note that penetrance values increase above this threshold more rapidly where the variance is greater, reflecting the broader pdfs for genotypes below the threshold value (see text for further discussion).

Given the intangibility of liability, this term might reasonably be replaced with *change in gene function*, as discussed above. Gene function may correspond to RNA or protein abundance at lower levels of gene function, or any other higher-level effect of a mutation on the casual path to a phenotype, such as a dynamical signal transduction readout within a tissue. In the same way that the classical definition of liability is the sum total of genetic and environmental risk factors for a particular phenotype, the penetrance value of a discrete phenotype would reflect the average change in gene function AND stochastic variation combined across *all* functional nodes within a network that converge on a phenotype. While liability, according to its classical definition, can be considered to be all genetic and environmental inputs into a system that define disease risk, the equivalent change in gene function is the highest-level readout(s) of biological function that causes a specific phenotype, and is the equivalent output derived from the input liability.

[We can see why we require such a high-level readout and its associated variation to capture all information about the genetic threshold value, and *vice versa*, as follows. A mutation that affects RNA abundance for instance will in turn influence protein abundance. As such, measurements of protein abundance will capture both the average mutational effect and associated noise at the RNA level as well as variation at the level of mRNA translation. In general, a higher-level readout will capture all of the mutational effects and variation at lower levels that feed into that particular node, as illustrated in Figure 2. Therefore, only a readout at a level that is ‘proximal’ to a discrete phenotype of interest will accurately reflect the genetic threshold value.]

**Figure 2.**
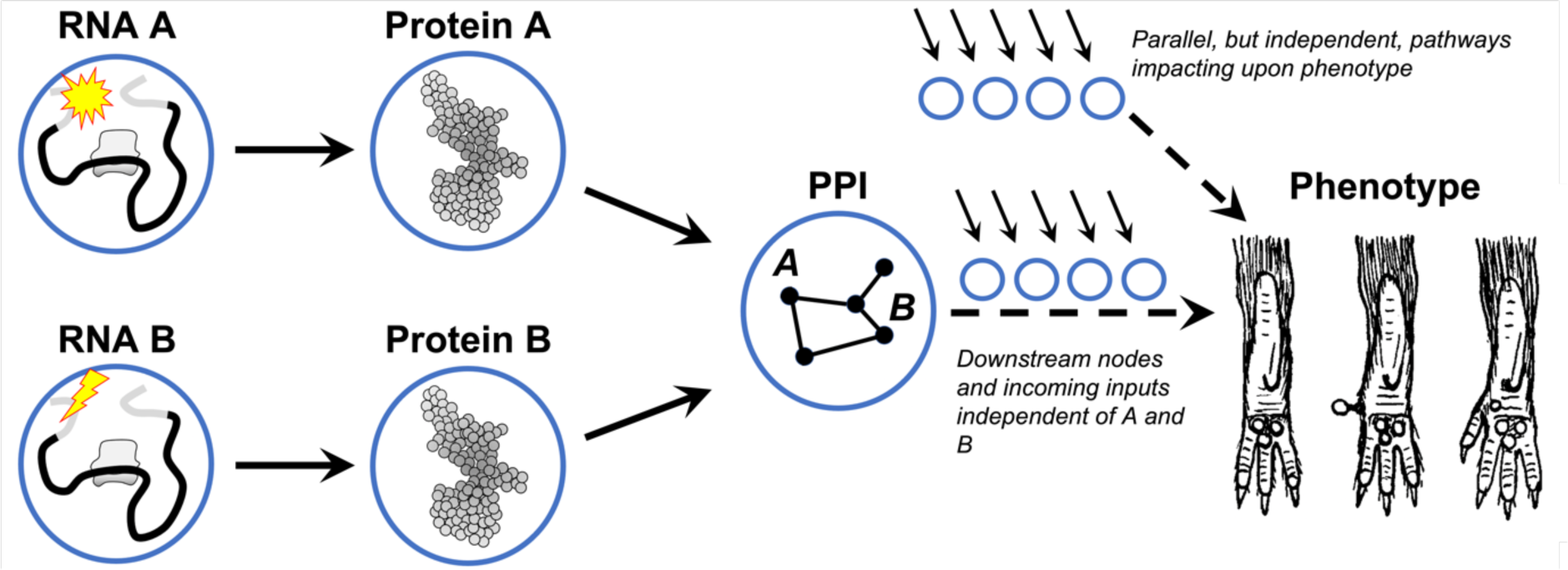
Relationships between mutations, pathways, networks and phenotypes. In the centre, we focus on a specific node – in this case a protein-protein interaction (PPI) complex – into which linear pathways relating to the synthesis of two components, A and B, feed (left). A reductionistic approach can be taken whereby the average mutational effects and associated variance in RNA, protein and PPI abundance and function can be measured. Downstream of *‘PPI*’ a series of steps eventually lead to a discrete phenotype. Although represented as linear, these components and functions form part of the wider network and the precise average values and stochastic variation associated with these components are influenced by these factors (arrows). By measuring ‘*PPI*’ we have no information about these values; however, the phenotype penetrance is influenced by all of these factors.

This high-level readout of gene function is the combined statistical distribution across all such nodes. In principle, this would allow for a parametric approach to calculate the genetic threshold value based on measurements of the relevant nodes within such a network. However, our limited understanding of the convoluted causal path linking functional readouts with phenotypes over time and biological scales precludes such an approach. The search for these predictive high-level readouts constitutes biomarker discovery, where a biomarker is a functional readout(s) that is either causally related to a phenotype or is correlated with a causal readout, and will be discussed further below.

In the absence of precise and detailed biological models such a parametric approach is not currently possible, and so the previous discussion of genetic threshold values was mainly illustrative. An alternative procedure that some researchers have used is to simultaneously score the penetrance of a discrete trait and to quantify a biochemical readout(s) for simple phenotypes that are apparent at the level of single cells. This removes the need for temporal separation of each assay and can be done at scale to include large numbers of individual cells collected in bulk. This simultaneous ascertainment approach removes the requirement for a detailed biological model linking readout and phenotype over time.

An example of this is an analysis of threshold cyclin-dependent kinase activity in the regulation of cell division during mitosis and cell division in *Schizosaccharomyces pombe* [43,44]. Multiple cyclin-CDK complexes regulate cell cycle progression in fission yeast, and earlier work had suggested that quantitative changes in total CDK activity led to the orderly initiation of S phase and mitosis. By using genetic simplification of the CDK network [59] in mutants expressing only a single cyclin-CDK chimera in place of the four cyclin-CDK complexes usually present, Swaffer et al. [43] demonstrated a progressive increase in a range of phosphorylation substrates until the end of the cell cycle. Initiation of phosphorylation of several substrates at different stages of the cell cycle was related to different cyclin-CDK affinities, suggesting that a causal relationship between CDK activity thresholds, differential substrate phosphorylation and initiation of G1-to-S and G2-to-M transitions.

To ascertain these threshold values, Patterson et al [44] developed a CDK activity biosensor which permitted the in vivo single-cell assessment of CDK activity and mitotic cell division. Fluorescence imaging allowed hundreds of individual cells to be simultaneously scored in bulk for cell division status, and their level of CDK activity to be quantified. By plotting CDK activity against the rate of division, a threshold of CDK could be seen directly. Below this threshold there was no cell division, but there was an exponential increase in the proportion of divided cells above this value. This is reminiscent of the exponential phase of a cumulative distribution function that relates stochastic variation to phenotype penetrance (Figure 1C).

A second study analysed threshold values within a gene regulatory network that specifies intestinal stem cell identities in *C. elegans* [13]. This network involves the maternal deposition of skn-1 transcripts that induce expression of end-1, which in turn activates expression of elt-2. There is also a parallel path in the network whereby skn-1 transcript regulates expression of elt-2 via other transcription factors also activated by skn-1 (Figure 3A). Using single molecule FISH, the authors were able to quantify processed transcript abundance for each of these components within intestinal cells, demonstrating a continuous distribution of stochastic variation in the expression levels of end-1 in *skn-1* mutants, ranging from complete loss to normal expression at the level of single cells. By contrast, end-1 expression was never zero in wild-types at the appropriate stage of development and demonstrated much less variability. Downstream of end-1, elt-2 demonstrated a bimodal distribution of expression in individual cells either falling within the normal range or with no detectable expression (Figure 3B.

**Figure 3.**
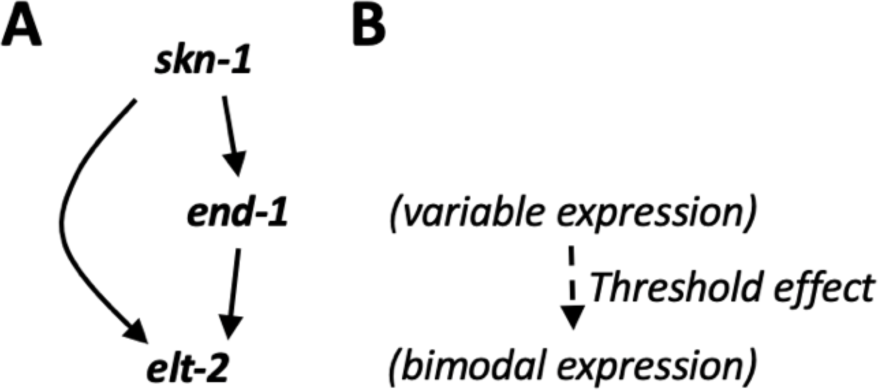
Simplified scheme of intestinal cell fate specification, adapted from [13].

To test the hypothesis that elt-2 expression only occurs if stochastic variation in end-1 transcript levels exceed a certain threshold value during a critical time-window, the levels of end-1 and elt-2 were simultaneously quantified in single cells. As for the CDK example outlined previously, this demonstrated a range of end-1 values for which no elt-2 expression was observed. A threshold value was defined as the maximum end-1 value above which elt-2 transcripts were non-zero in simultaneous quantification (Figure 3B).

These studies offer additional layers of insight into threshold effects. In the CDK study, the same analysis was performed on both a wild-type and a CDK mutant strain, which suggested that the threshold value was the same for both. However, once the threshold had been reached the phenotype penetrance (probability of cell division) increased more sharply with increasing CDK activity in the mutant versus wild-type strain. This is consistent with a cumulative distribution function representing a greater level of stochastic variation in the mutant. In the intestinal specification study, the threshold value for two high penetrance *skn-1* alleles were the same, whereas for a weaker mutation, the threshold was lower. In this case it was suggested that skn-1 may partially activate the parallel pathway, thereby changing the topology of the network and redefining the threshold for end-1 activity. In this case, ascertainment of threshold values therefore had the added advantage of revealing a systems level property of the gene regulatory network.

This leads to two fundamental aspects of stochasticity. Firstly, while the average effects on gene function may differ for different genotypes, threshold values are properties of wider network topology: they are independent of genotype and remain constant for a given genetic background. In dynamical systems modelling, similar emergent properties of signalling pathways have been uncovered including distributed robustness and bifurcation points that determine cell fate decisions [56,57]. Second, stochastic noise is almost always greater in networks carrying a defined mutation (or any other specific perturbation) (Figure 1C). This may relate to buffering mechanisms and localised network properties that confer robustness.

[This note addresses two points that were raised in review. It was stated in the previous paragraph that the threshold value is invariant for a given genetic background. This requires some justification as Wright does indeed assume constant threshold values across genetic backgrounds [31,32]. In Waddington’s view [64], the epigenetic landscape constitutes the wider network consisting of the biophysical parameters that define the nodes and edges within a dynamical system (such as a large collection of ordinary differential equations). If the threshold value is a property of this wider network, rather than the mechanisms that modulate effects of a specific mutation (i.e. proximal to the mutation), then it would be invariant for a given genetic background in a controlled environment. It is also important to note that the threshold value is not ‘picked’ arbitrarily (i.e. scaled and shifted appropriately to reflect both the liability values and the penetrance of the mutation). Rather, it reflects network properties and is not peculiar to a particular mutation].

### Could genetic threshold values be determined for complex (morphological) phenotypes?

These examples of simultaneous ascertainment demonstrate the possibility of determining threshold values for relatively simple cellular phenotypes. Could this be possible for complex phenotypes at the level of whole tissues occurring late in the process of development? It was stated above that a molecularly targeted genetic threshold model requires assessment of a combination of nodes that capture the total stochastic variation within the wider network that impacts upon the phenotype. A potential pushback on this idea is that this is an unknowable causal graph, and the strictly sufficient measurements (e.g. every protein molecule concentration, modification state, and location in every cell) are unrealistic to obtain without some strong casual assumptions. Is such a high-level readout that captures all relevant functional variation just as intractable as the classical definition of liability?

A great deal of work will be necessary to answer this question definitively, but it is illustrative to consider a practical scheme using single cell RNA-sequencing (scRNAseq) and spatial transcriptomic technologies. In his epigenetic landscape model, Waddington [64] proposed that only a limited set of discrete cellular and tissue-level fates can be arrived at over the course of developmental time in an individual animal (i.e. amongst groups of genetically identical cells). Each path within the landscape reflects its tolerance to stochastic variation and resistance to changes in cell fate. His idea also encapsulated tissue transplantations, thereby traversing all tissues within an embryo and reflecting the concepts of specification and determination in experimental embryology. As well as his famous epigenetic landscape diagram, Waddington also considered a ‘phase-space box diagram’ consisting of three dimensions. One dimension was time, and the other two functioned to define clusters of cellular identities. These clusters overlapped and each identity mapped onto future clusters, representing specification and differentiaton over time.

While this diagram was entirely theoretical and heuristic at the time, it bears remarkable similarity to what we are very familiar with from scRNAseq datasets where different cell types are revealed as clusters following dimensionality reduction. scRNAseq is necessarily destructive of the samples analysed and thereby strictly prohibiting temporal analyses (althoug trajectory analyses are possible given certain assumptions). However, it is possible to conceive of a simultaneous ascertainment approach, firstly using clustering to define discrete cellular phenotypes at the single cell level, and then using a form of principal component analysis to define quantitative stochastic variation that correlates with these cell types. In essence, this would serve to define both the high-level quantitative readout and associated variation that was discussed reviously and suggests that this form of liability is ascertaniable. In terms of higher-level phenotypes, more ambitious and exploratory models that relate these single cell phenotypes within a tissue to the penetrance of higher-level phenotypes would be required. Possible models include the number of each cell type (total/differentiated/proliferative..) with or without weightings according to location or developmental time. Alternative models could be scrutinised by evaluating a variety of interventions using do-calculus.

While speculative, this illustrates how such a molecularly target liability model based on stochasticity might be possible.

### Comparing experimental approaches to threshold determination with causal inference

The simultaneous ascertainment of thresholds described above is based on a direct comparison of the values for a particular readout of gene function in cells with or without a phenotype. Both ‘affected’ and ‘unaffected’ cells come from a mixed group that demonstrate stochastic variation in the values of the readout and incomplete penetrance for the trait. This approach is based on observational data alone. The stochasticity is useful in generating a distribution of values for a biological variable that would be difficult to achieve experimentally without artefact. It also captures natural biological variation.

Controlled experiment is the gold-standard for demonstrating causality. This is because correlations between parameters could arise from covariance of two readouts with a third confounding variable and not only through a causal relationship between them. In analysing heterogeneous human populations, it is not possible to control for all confounders, and in this case randomised control trials (RCTs; also invented by Fisher) are the gold-standard for establishing causality. Here, confounders are controlled for by *random assignment* of patients to different treatment groups. In analogy, stochasticity can also be used to achieve a form of random assignment to infer causal relationships.

In genetics, the stochastic nature of chromosomal segregation and crossing over provides a form of randomisation by which causation can be inferred. Linkage analysis has been used to identify thousands of Mendelian disease genes [1-4,45,46]. The premise for this analytical approach is to first establish the mode of inheritance for a particular trait (e.g. dominant or recessive) and to genotype polymorphic genetic markers throughout the genome. Segregation and independent assortment of these variants occurs randomly with respect to disease status in regions of the genome that are not linked to the disease such that a particular autosomal allele has a 50% chance of passing from a parent to their offspring. As in RCTs, Mendelian inheritance serves as randomisation with precisely defined ratios for the assignment of different regions of the genome to case (affected) and control (unaffected) groups. Deviation from this baseline value indicates linkage to a causative mutation. A special form of this causal analysis for quantitative traits is the transmission-disequilibrium test [47–50]. Another major area of causal inference is known as Mendelian randomisation whereby genetic susceptibility variants for a specific phenotype are used as *instrumental variables*, and control for confounding of a risk factor under study [51,52]. Observational data with randomisation through stochastic processes therefore allows causal mechanisms to be inferred.

We saw from the previous examples of threshold determination through simultaneous assessment of single cells that stochastic variation in biological networks can provide causal information. Specifically, for a mutant exhibiting a discrete phenotype, stochastic variation in a causative readout will exhibit a discontinuous correlation with this phenotype i.e. there will be a correlation only above a threshold value relating to the truncated distribution in excess of the threshold (i.e. the line above blue shading in Figure 1A). Furthermore, while the correlation coefficient may differ between strains with different mutations, the threshold value is constant for a particular isogenic strain where a high-level functional liability readout is quantified. Here, the randomisation afforded by stochastic processes together with the concept of an invariant threshold value for a discrete trait and a uniform genetic background could provide a means to infer causal relationships in complex biological datasets. This concept of discontinuous correlation has parallels with an approach that is frequently used in the social sciences and econometrics known as regression discontinuity (RD) [53].

### Structural equation models in causal inference and stochasticity

Causal inference (CI) is a statistical approach which allows sources of variation within a causal model to be quantified in relation to a variable outcome [54,55]. The causal model will have been derived from experimental evidence and/or will constitute a hypothesis relating to causation. As such, causal inference can only be applied relatively late within the discovery timeline of a research question, whereby extensive prior research has established high confidence causal relationships between various components. As will be described, it has several uses. CI can accommodate unknown confounders thereby allowing incomplete biological models to be analysed. It can quantify the relative contribution of different sources of variation to variable (phenotypic) outcomes in an unbiased way, and it can be used to infer these values even for variables that cannot be measured directly.

In CI a causal model is represented by a structural equation model (SEM) whereby nodes represent components within the system and edges are represented as arrows that depict causal relationships between these components. The purpose of CI is then to calculate weights for these edges that correspond to the proportion of variation in a particular outcome (such as a phenotype) that is accounted for by variation in a particular node. The weights for each edge are determined by calculating correlations between the values of different nodes (Figure 4).

**Figure 4.**
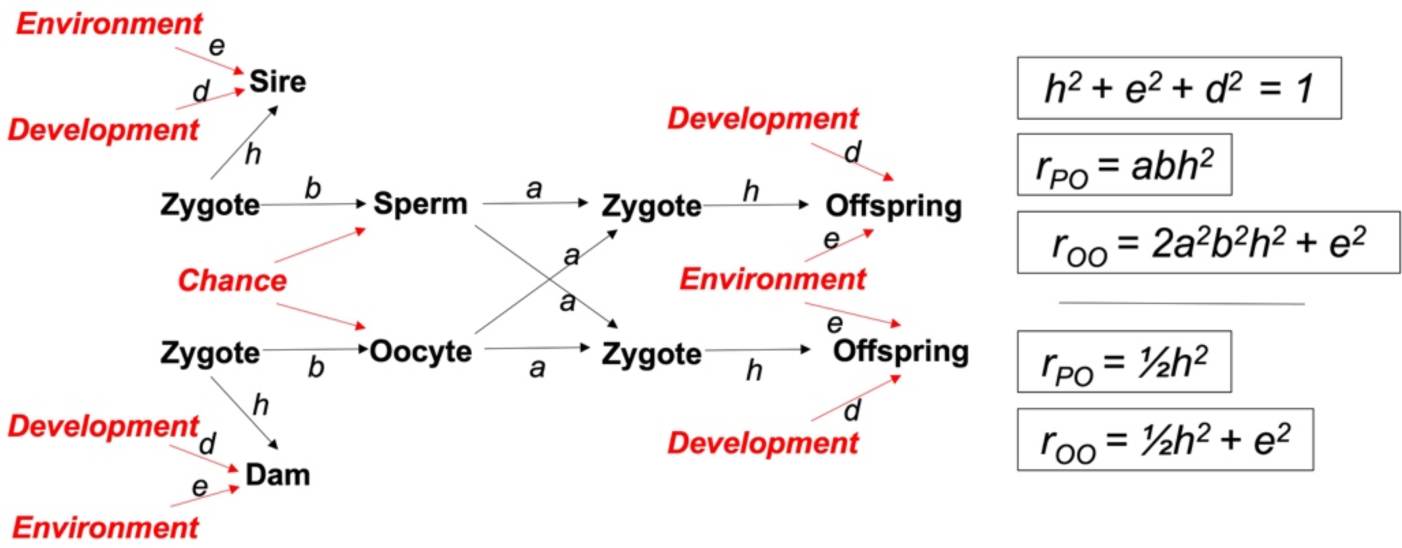
Structural equation model showing paths linking parental (sire, dam) phenotypes with offspring phenotypes. Nodes are capitalised: tangible entities are in black, unmeasurable entities in red. Pathway coefficients are given in lower case (*a,b,e,d,h*). Simple mathematical relationships linking pathway coefficients with linear correlation coefficients between quantitative parent-offspring and offspring-offspring phenotypes are listed. With basic algebra, *h,e,d* can be derived although these would not otherwise be quantifiable. (Adapted from [60]). Note, Wright referred to ‘*Chance’* as irregularities of development and also refers to ‘..ontogenetic irregularity..’ in relation to the term *D,* hereby labelled *Development*. *d* might reasonably represent weighting for stochastic processes, discussed in the text.

A feature of SEMs that are amenable to CI is that they are directed acyclic graphs (DAGs). This means that there is a path of causation following a series of nodes to a specified outcome without forming a closed loop. This form of pathway analysis was first proposed by Wright in 1920 [60], where he applied it to breeding experiments that he had performed in various strains of guinea pigs that demonstrated differences in coat colouration, and later generalised in 1921 [58]. By formulating a scoring system to quantify this variation and normalising the data, he was able to calculate the linear correlations between coat colouration in parents and their offspring, and between offspring within the same litter. He considered an SEM whereby fertilised zygotes gave rise to each of the parents’ coat colouration phenotype as well as the parents’ gametes (Figure 4). Through the precise statistical ratios of Mendelian inheritance, the gametes from each parent combined to generate the zygotes that generated the subsequent generation. In both generations, the observed coat colour phenotypes of both the parents and the offspring were defined by genetic and environmental variables, as well as a third term which Wright coined ‘*Development’* to account for the ‘ontogenetic irregularities’ i.e. stochastic processes.

By incorporating correlations between parents and siblings, and between siblings within the same litter into the SEM, it was possible to quantify the contribution of genetics, environment and development to variation in phenotypic outcomes. In this way this statistical approach was able to infer and quantify the contribution of variation in these otherwise intractable parameters to variation in coat colour. As a form of sanity check, Wright showed that environmental variation made the same contribution to phenotypic variability in both an isogenic line and an outbred strain of guinea pigs. By contrast, his calculation showed that genetic variation contributed significantly to phenotypic variability in the outbred strain but made no contribution to variability in the isogenic line.

Since Wright, CI was largely forgotten in the biomedical sciences, especially because of work by his contemporaries such as Galton/Pearson and Fisher who developed statistical methods for linear regression in genetics and randomised control trials (RCT), respectively. They established the mantra ‘correlation not causation’ [reviewed in 61]. They asserted that causal relationships could not be deduced from correlations within population-based datasets and could only be derived from RCTs, and this notion predominated. However, Wright had shown that correlations could reflect causation where a high confidence SEM is available.

CI based on SEMs has undergone extensive development. As stated by Meinshausen et al. [62], ‘an SEM consists of: (a) an underlying true causal influence diagram for random variables that are represented by nodes within the DAG, and; (b) a function that relates each variable to their parental variables and an error term. This so-called ‘*do-operator’* sets a particular variable to a deterministic value according to the SEM that relates the variable to its parental node, and can be applied to several variables simultaneously. This is a conditional probability of the kind – ‘what is the probability of a specified outcome given that I assign a particular value to a specified variable (i.e. an intervention)’’ [54,62,63].

What might the utility of CI in understanding stochastic processes be? It might seem that, if a causal SEM is known, then there is no need for CI. However, CI would allow for a causal relationship to be defined in an SEM that subsequently turns out to have a weighting of zero. In other words, it may turn out that a parameter that exhibits some degree of variability and is thought to contribute variation to a phenotypic outcome might actually make no contribution at all. I began this *Perspective* by defining different sources of stochastic variation. Another utility of CI could be to apply weights to these different sources of variation, thereby highlighting the importance of one or other form of stochastic variation associated with *a particular combination of mutation*, *genetic background and environment.* The complexities of the network that constitutes an SEM may unexpectedly render the system robust to variation in a particular node. In dynamical systems modelling, similar emergent properties of signalling pathways have been uncovered including distributed robustness and bifurcation points that determine cell fate decisions [56,57].

### Conclusions and perspectives

In this *Perspective*, I have considered how stochastic processes could provide insight into mechanisms of pathogenesis. In isogenic model systems, stochastic variation is the major source of phenotypic variability and provides information about genetic threshold values where incomplete penetrance is observed. This is because there is an exact parametric relationship between phenotype penetrance and the probability distribution of stochastic variation. However, in order to make such a calculation, the sum total of average mutational effects and stochastic variation must be quantified across all nodes within a biological network that converge on a trait of interest. This has been achieved for relatively simple phenotypes by simultaneous quantification of a continuous biological readout and phenotype penetrance. However, this is unlikely to be possible for complex morphological traits where mechanisms of pathogenesis for such high-level traits are not well understood.

Causal inference may provide an alternative means to harness this information that is afforded by stochastic processes. It may be possible to determine the degree to which different sources of stochastic variation within a network contribute to overall phenotypic variability. The main utility of CI is to quantify the degree to which different sources of stochastic variation within a network contribute to overall phenotypic variability. This approach focuses on high-confidence causal relationships established through controlled experiments whereby SEMs can be drawn. An advantage of CI is that large gaps in the causal diagram can be tolerated by treating them as unknown confounders, and so a partial analysis of known causal mechanisms could be undertaken. By calculating correlations between different nodes within such a model, the extent to which variation in each node contributes to overall phenotypic variability can be estimated. It also allows inferences to be made for nodes that can’t be measured (but which are known) through these indirect calculations. In this way, unexpected and non-trivial weights can be given to known components in a causal framework.

This could help to disregard molecular and cellular mechanisms that seem intuitively to be causally related to a phenotype, potentially revealing novel mechanisms of buffering and robustness. It could also help to prioritise molecular targets for therapy. Genetic therapies are designed to target high-confidence causal mechanisms within the central dogma of molecular biology and many mutations affect multiple modes of gene function (RNA, protein etc). Weighting the relative contributions of different modes of gene function to phenotypic variability may therefore help to prioritise therapeutic targets. By making precise statistical statements for different causal relationships, CI may allow quantitative statements regarding the effectiveness of a particular therapy to be made. Similarly, it may permit the design of adjunct or combined therapies, targeted to a particular mode, where residual disease risk is present. In future, it may also be possible to make quantitative predictions about disease outcomes for individual patients undergoing a particular therapy. This would require disease models that represent a patient’s total genetic constitution in which disease outcomes can be predicted (e.g. organoids, assembloids etc).

## Acknowledgements

This work was funded by a Wellcome Trust Collaborative Award in Science (210585/Z/18/Z) to DJ. This research was also supported by the National Institute for Health Research Biomedical Research Centre at Great Ormond Street Hospital for Children NHS Foundation Trust and University College London.

## Notes

### Competing Interest Statement

The authors have declared no competing interest.

### Summary of Updates

Corrected minor errors and inserted additional information relating to epigenetic landscapes.

